# Single cell analysis of population-wide nuclear and cytosolic drug responses using high-content FRET imaging: measuring protein kinase activation in rat primary striatal neurons

**DOI:** 10.1101/2021.01.31.429059

**Authors:** Jace Jones-Tabah, Ryan D. Martin, Jason C. Tanny, Paul B.S. Clarke, Terence E. Hébert

## Abstract

Genetically-encoded biosensors are used to track biochemical activities in living cells by measuring changes in fluorescence emitted by one or more fluorescent proteins. In the present article, we describe the application of genetically-encoded FRET biosensors with high content microscopy to image the signaling responses of thousands of neurons in response to drug treatments. We applied this approach to reveal intercellular variation in signaling responses among cultured striatal neurons stimulated with multiple drugs. The striatum is largely composed of medium-spiny GABAergic neurons which are divided into two broad sub-types based in part on their expression of dopamine D1 *vs.* D2 receptors. Using high content FRET imaging and immunofluorescence, we identified neuronal sub-populations with unique responses to pharmacological manipulation. Focusing on dopamine- and glutamate-regulated PKA and ERK1/2 signaling in both the cytoplasm and nucleus, we identified pronounced intercellular differences, in both the magnitude and kinetics of signaling responses to drug application. Importantly, we found that a conventional “bulk” analysis that included all cells in culture yielded a different rank order of drug potency than that revealed by our single-cell analysis. The high degree of heterogeneity that we observed at the single cell level would not have been detectable using common population-level analyses, derived for example from western blotting or plate reader-based measurements. In conclusion, our single-cell analytical approach highlights the limitations of population-level analyses, and provides a novel way to study signaling biology.

## Introduction

Genetically-encoded biosensors based on fluorescent proteins allow researchers to track the biochemical activity of living cells with imaging technologies that detect changes in fluorescence emitted by one or more fluorescent proteins. Many such biosensors are based on the principle of fluorescence resonance-energy transfer (FRET) ^1^. FRET biosensors have been constructed to report many signaling processes, including changes in the concentration of ions and small molecules ^2,3^, enzymatic activities of protein kinases and GTPases ^4,5^, and the dynamics of post-translational modifications such as phosphorylation and acetylation ^6^. While there are many ways to measure FRET, fluorescent plate-readers and fluorescent microscopes are the most commonly used, and each offers specific advantages. Plate readers provide high-throughput but lack cellular resolution. In contrast, conventional microscopes provide high spatial and temporal resolution but provide only low throughput, and time-resolved imaging of large numbers of cells presents a challenge. An alternative approach, high content microscopy, provides the benefits of cellular resolution and the ability to image many cells in multi-well formats. Although lacking the sub-second temporal resolution of conventional microscopy, automation allows for time-resolved imaging to be performed across many conditions with minimal experimenter involvement. In this article, we describe the application of high content imaging with genetically-encoded FRET biosensors to study responses to drug stimulation at the level of individual cells. Specifically, we examine the activation of protein kinase A (PKA) and extracellular regulated kinases (ERK1/2) in the cytosol and nuclei of primary striatal neurons following exposure to multiple drugs. Although we focus on the application of FRET-based biosensors, the pertinent analytic principles can be applied to any fluorescent indicator.

Neurons in the dorsal striatum regulate goal-directed locomotion, behavioral action selection, and specific forms of learning. Medium-spiny GABAergic projection neurons (MSNs) account for 95% of the neurons found in the rodent striatum ^7,8^. These neurons integrate dopaminergic and glutamatergic inputs to the striatum and send inhibitory projections to several downstream nuclei. MSNs are divided into two populations, based on both anatomical and molecular features: “direct pathway” MSNs, which express the Gα_s/olf_-coupled D1 dopamine receptor (D1R), and “indirect pathway” MSNs, which express the Gα_i/o_-coupled D2 receptor (D2R) ^9^. Within these broad neuronal subtypes, there is also variability. Recent studies have used single cell transcriptomics to demonstrate that within striatal subtypes, there is heterogeneity of molecular identities ^10–13^; however, the functional consequences of this transcriptional diversity remain unexplored. Both MSN subtypes also express NMDA-sensitive glutamate receptors, which are involved in regulating plasticity at cortico-striatal synapses. In direct pathway MSNs, D1R and NMDA glutamate receptors cooperatively regulate synaptic plasticity and gene expression through activation of PKA and ERK1/2 kinases ^14–16^, and activation of these intracellular signaling pathways plays an important role in the effects of cocaine ^17,18^. However, in a recent study using single cell transcriptomics, only a small proportion of molecularly-defined direct-pathway MSNs were found to respond to cocaine *in vivo* ^19^. It is unknown how much of this *in vivo* variation can be attributed to differences at the circuit level, to intrinsic variability in the properties of individual cells, or to both. We therefore sought to determine whether the signaling responses of cultured striatal neurons to dopaminergic and glutamatergic stimulation would also show intercellular heterogeneity. To this end, we used genetically-encoded FRET biosensors of PKA and ERK1/2 activity, in combination with high content confocal microscopy, and measured signaling responses simultaneously from thousands of individual striatal neurons in culture. By complementing FRET measurements with morphological analysis and quantitative immunofluorescence, we identified neuronal sub-populations with unique responses to pharmacological manipulation.

## Materials and Methods

### Drugs and Reagents

Unless otherwise noted, products were purchased from Sigma-Aldrich. SKF 81297 hydrobromide (Toronto Research Chemicals), forskolin, U0126 and SCH 772984 (Selleckem) stocks were prepared at 10 μm in DMSO and stored at −80°C. *N*-Methyl-D-aspartic acid was prepared fresh in HBSS.

### Animals

Sprague-Dawley dams with postnatal day 1 pups were purchased from Charles River, Saint-Constant QC, Canada. Animals were maintained on a 12/12-hour light/dark cycle with free access to food and water. All procedures were approved by the McGill University Animal Care Committee, in accordance with Canadian Council on Animal Care Guidelines.

### Isolation and culture of primary striatal neurons

Primary striatal neurons were prepared from postnatal day 1 pups as previously described ^20^. One day in advance, 96-well optical bottom imaging plates (Nunc) were coated overnight with 0.1 mg/ml poly-D-lysine dissolved in phosphate buffered saline (PBS for cell culture - Sigma). Prior to dissection, plates were washed three times with sterile water and allowed to dry, and all required solutions were warmed to 37°C. Pups were decapitated and brains were rapidly removed and placed in ice cold Hank’s balanced salt solution (HBSS) without calcium and magnesium (Wisent). Striata were dissected, and then digested on a rotator at 37°C for 18 min with papain diluted to a final concentration of 20 units/ml in a neuronal medium (Hibernate A Minus Calcium, BrainBits). Next, in order to halt digestion, tissue was transferred to HBSS containing 10% fetal bovine serum, 12 mM MgSO_4_, and 10 units/ml DNase1 (Roche) for 2 min. Striata were pelleted by centrifugation at 300 X *g* for 5 min and supernatant was removed. Tissue was then triturated in HBSS with 12 mM MgSO_4_ and 10 units/ml DNase1 using a fire-polished Pasteur pipette. Once a cell suspension was produced (approximately 20 up and down triturations), it was passed through a 40 μm mesh filter (Fisher) to remove undigested tissue and then centrifuged on an OptiPrep™ gradient as previously described ^21^, in order to remove cell debris. Purified neurons were then counted and diluted in Neurobasal™-A Medium (NBA) with 1X final concentration of B27 supplement (Gibco), 1% GlutaMAX (Gibco) and 1% penicillin/streptomycin (henceforth referred to as complete NBA) supplemented with 10% fetal bovine serum. On a pre-coated 96-well plate (see above), 50,000 cells were plated in 75 μl volume per well. Sixteen hours after plating, cells were washed with HBSS without calcium and magnesium, and media was changed for complete NBA, containing 5 μM cytosine-D-arabinoside to inhibit cell proliferation. Cultures were maintained in complete NBA, and media was refreshed by exchanging 30% of the volume with fresh media every 3 days.

### Virus production and transduction of primary neurons

The FRET-based protein kinase biosensors AKAR3-EV and EKAR-EV were generously provided by Dr. Michiyuki Matsuda ^4^ and expressed with either nuclear export (NES) or nuclear localization (NLS) peptide sequences. FRET biosensors were expressed using a neuron-specific adeno-associated virus plasmid, pAAV-SynTetOFF ^22^, kindly provided by Dr. Hiroyuki Hioki. All experiments were performed with AAV serotype 1 produced by the Neurophotonics Platform Viral Vector Core at Laval University. Primary striatal neurons were transduced by adding AAV directly into the culture media three days after cell plating, using a multiplicity of infection of 5000 viral genomes/cell. Neurons were then maintained as described above for 7 days prior to imaging.

### High content FRET imaging in primary neurons

Live-cell imaging was performed at 37°C and with 3% CO_2_ using an Opera Phenix™ high content confocal microscopy system (Perkin Elmer). One hour prior to imaging, media was replaced with 90 μl of HBSS with calcium, magnesium and sodium bicarbonate (Wisent). For antagonist experiments, antagonist or vehicle was also added at the appropriate concentration. Plates were transferred to the Opera Phenix and allowed to acclimatize in the live-cell chamber for 10 min before baseline image acquisition. Images were acquired using a 40X water-immersion objective using a 425 nm laser for excitation of CFP. Emissions were detected with filters at 435-515 nm (CFP) and 500-550 nm (YFP). For time-course experiments, approximately 20 fields were imaged per well, with fields being evenly spaced across the well. Baseline images were acquired and then vehicle or drug solution was added directly to each well using a multichannel pipette. Drugs were added at a volume of 10 μl to achieve the required final concentration in the well. Images were then acquired at 10-minute intervals for a total of 50 min. Following each imaging session, cells were fixed and could be processed for immunofluorescence, as described next.

### Cell fixation and immunofluorescence

Cells were fixed for 10 min in 2% paraformaldehyde prepared in PBS. Fixed cells were then washed twice with PBS and permeabilized for 10 min using 0.3% Triton X-100 in PBS. Blocking was performed for 3 hours at 4°C with 5% bovine serum albumin (BSA) in PBS. Next, cells were incubated overnight at 4°C with primary antibodies diluted in PBS with 5% BSA, then washed twice with PBS, and subsequently incubated for 3 hours at room temperature with secondary antibodies diluted in PBS with 5% BSA. After two additional washes with PBS, cells were incubated with Hoechst dye (Invitrogen) diluted 1:10,000 in PBS. The following primary antibodies and dilutions were used: anti-DARPP-32 (1:1000, catalog #2302, Cell Signaling Technology), anti-cFos (1:2000, catalog #sc-52, Lot #C1010, Santa Cruz) and anti-beta tubulin (1:4000, catalog #801213, Lot #B272898, BioLegend) Secondary antibodies were Alexa 488 anti-mouse (1:1000; A21236) and Alexa 647 anti-rabbit (1:1000; A21245), both from Invitrogen. Fluorescence imaging of fixed cells was performed in an Opera Phenix^™^ high content confocal microscope at 40X magnification.

### Image processing and fluorescence quantification

All image analysis was performed using Columbus™ analysis software (Perkin Elmer) using the following generalized workflow. For FRET imaging, transduced cells were automatically identified using “Method B” run on the combined CFP and YFP fluorescence intensity. Specific thresholds for object seize, brightness and contrast were optimized for each sensor and then kept constant for all experiments. The identified objects were then further filtered based on morphology and fluorescence intensity to exclude dead cells, non-cell objects and clusters of overlapping cells. Specific parameters were determined by visual inspection and were adjusted for each sensor. After filtering the object population, CFP and YFP intensities at each pixel were used to calculate the FRET ratio (YFP/CFP), which was then averaged to create a single FRET ratio for each object at each time-point. Values for fluorescence intensities, FRET ratio, object area, object roundness and spatial coordinates within the image were calculated and exported as text files for further analysis. The same pipeline was applied for quantitative immunofluorescence experiments, except that nuclei were first detected using the Hoechst stain, and then this was used as a mask to define a region in which fluorescence signals from the Alexa-488 and Alexa-647 channels were then quantified.

### Single cell analysis

For time-course FRET imaging, single cells were tracked across time points using an analysis script written in R (available upon request) and applied to the output of the image analysis. Objects were matched between time-points using the Cartesian coordinates of each object in the image, with a threshold for maximum displacement between subsequent images. To ensure correct matching of objects at each time-point, the size and fluorescence intensity of each object was cross-checked, and objects which exhibited a change in size or brightness of greater than 30% were excluded from analysis. Only objects that could be positively matched across all 6 time points in the experiment were included for analysis. ΔFRET values were then calculated for each object relative to the object’s baseline FRET, and converted into percent changes in FRET (%ΔF/F). Here, the denominator (F) was calculated as the average baseline FRET, not of the particular cell in question, but rather of *all* cells sampled in the same well. This approach avoided extremely high %ΔF/F values that tended to occur with cells having a very low baseline FRET value.

Values corresponding to single cells from independent experiments were then pooled for data visualization in heat maps. Grouping of FRET responses by either magnitude or kinetics was performed by time-series clustering using the TSclust package in R using the “pam” and “shape” functions for clustering by response magnitude and kinetics, respectively ^23^. All figures were generated as R markdown files. R scripts for analysis and data visualization are available upon request.

### Statistical analysis

All statistical testing was performed in R using the rstatix package. Data in **Fig. 1** and **Fig. 5** were analyzed using Bonferroni-corrected t-tests, with each treatment being compared to DMSO. In **Fig. 2** data were analyzed first by 2-way ANOVA, using treatment and cell-type as factors. ANOVA was followed by multiple comparisons performed using Bonferroni-corrected t-tests, comparing “Medium Round” cells to all other cell-types.

**Figure 1:**
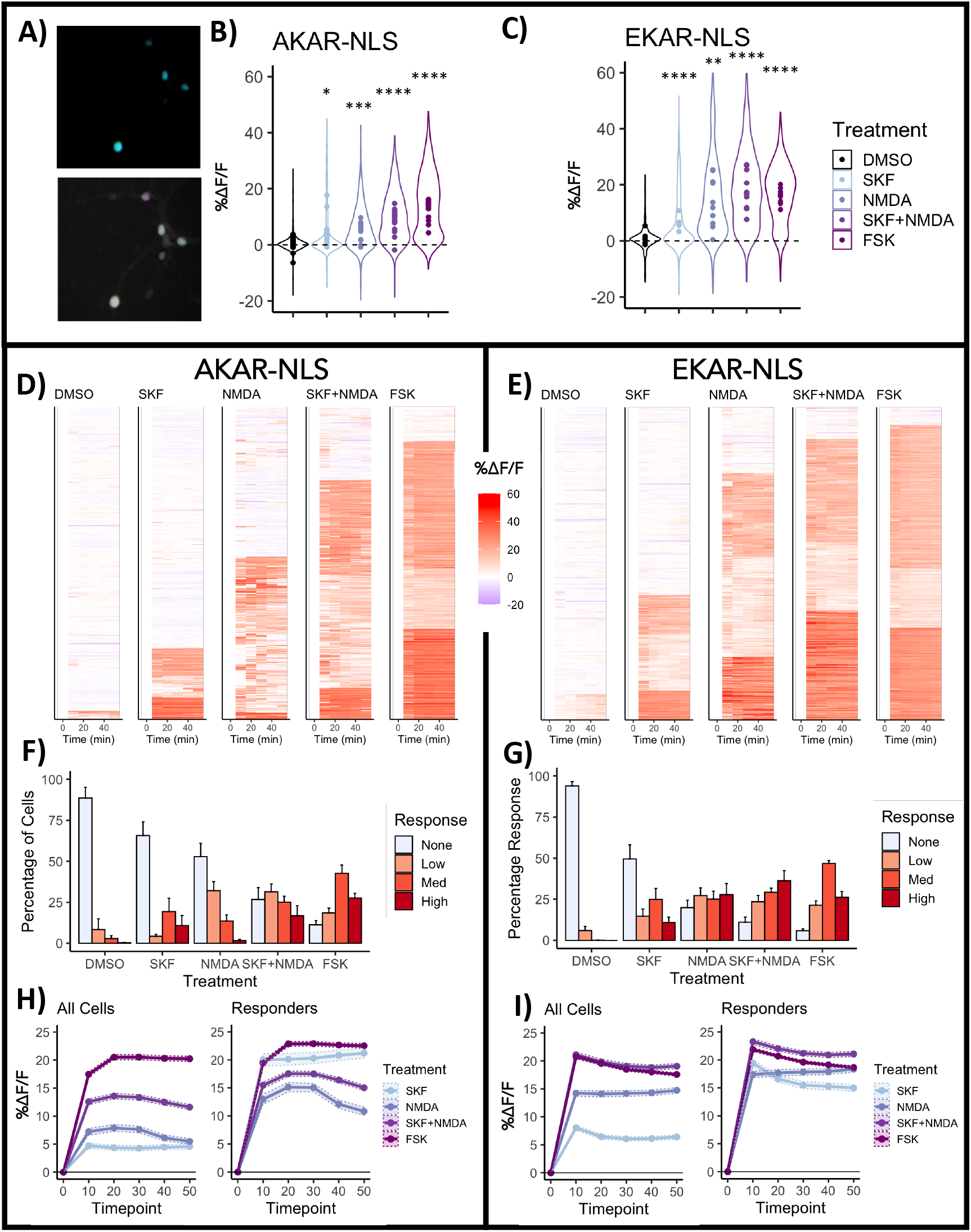
High-content imaging of nuclear PKA and ERK1/2 activity with FRET biosensors. A) Representative image of primary striatal neurons expressing a FRET reporter tagged with nuclear localization signal (NLS) (top) and outlines showing object identification by Columbus software B, C) Percent change in FRET ratio (%ΔF/F) 10 min after drug stimulation. Violin plots show the range of *individual* cell responses, while points indicate the mean of each *biological replicate*. D, E) Heatmaps depicting individual cells imaged at 10-minute intervals. Cells are organized into the four response clusters described in the following panels. F, G) Percentage of cells falling into each of four response clusters (mean±SEM across biological replicates). H, I) Comparison of the mean %ΔF/F time-course for each drug when all cells are included (left) or after removal of non-responding cells (“None” cluster) (right). Shaded region shows the SEM calculated across cells. Experiments consisted of 10 and 8 biological replicates for AKAR-NLS and EKAR-NLS respectively. Statistical comparisons were performed by Bonferroni corrected t-tests comparing each treatment to DMSO. *p<0.05, **p<0.01, ***p<0.001, ****p<0.0001. Abbreviations and drug concentrations: DMSO, dimethyl-sulfoxide; SKF, SKF 81297 (1 μM); NMDA, N-methyl-D-aspartic acid (5 μM); and FSK, forskolin (5 μM).

**Figure 2:**
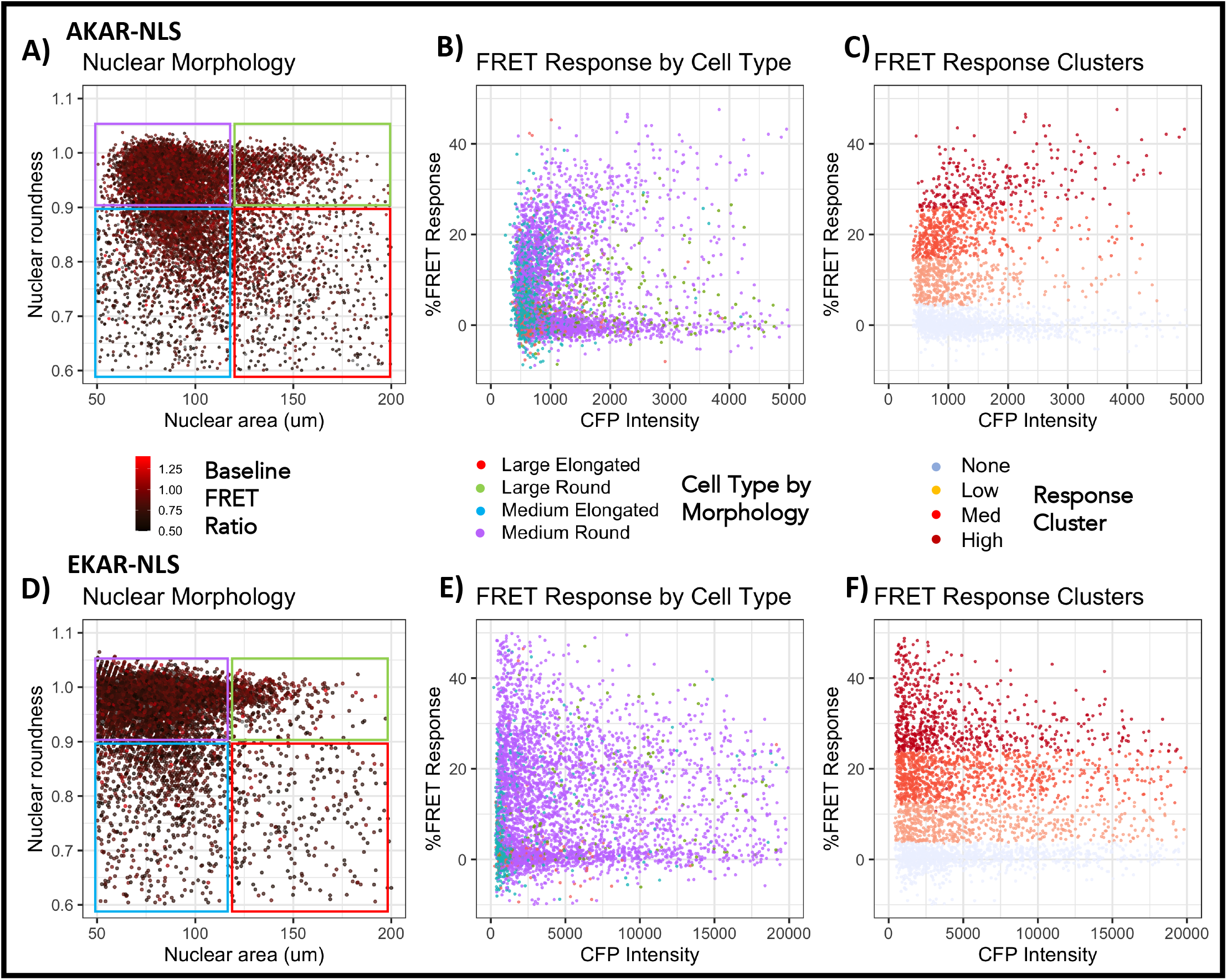
Assessing basal FRET ratios, cellular morphology, FRET responses and biosensor expression levels. A,D) Baseline FRET ratio (point color) of single cells, as a function of nuclear size and shape (i.e. cross-sectional area and roundness, respectively). Based on nuclear size and shape, cells were individually assigned to one of four categories, framed by the colored rectangles. B,E) Relationship between FRET response (%ΔF/F) and biosensor expression level (CFP intensity, arbitrary units) across morphology-defined cell types. C,F) Relationship between FRET response (%ΔF/F) and biosensor expression level (CFP intensity, arbitrary units) within “medium round” cells. Colors indicate the four response clusters (i.e. none, low, medium, and high, as described in **Figure 1)**. For all panels, data are merged from all treatment conditions.

## Results

In the experiment presented in **Fig. 1**, we expressed nuclear-localized reporters of PKA (AKAR-NLS) and ERK1/2 (EKAR-NLS) in primary striatal neurons and we performed live-cell high content imaging (**Fig. 1A**). Neuronal cultures were treated with either vehicle, SKF 81297 (a selective agonist of D1-like receptors), *N*-methyl-D-aspartic acid (NMDA, an agonist of NMDA-type glutamate receptors), a combination of these two drugs, or forskolin (a direct activator of adenylyl cyclase). **Fig. 1B-C** displays the FRET changes of individual cells (violin plot) and biological replicates (points) after a 10 minute exposure to the specified drugs. While NMDA receptors are ubiquitously expressed in all striatal MSNs, D1Rs are expressed only in dMSNs ^8,9^, and thus even assuming a pure population of striatal MSNs, should be present in no more than ~50% of cultured neurons. Single cell analysis allowed us to visualize the responses of individual cells over the imaging period (**Fig. 1D-E**). To better visualize differences between individual cells, the responses to all drugs were pooled and sorted into four groups using the magnitude of FRET change as the primary clustering variable. This clustering produced four response-types which we defined as non-responsive cells, low, medium and high responding cells, based on the average response magnitude of the cluster. Within each response-type, we were able to analyze drug responses both in terms of average response magnitude of biological replicates (**Fig. 1B-C**), and percentage of responding cells in each replicate (**Fig. 1F-G**). This revealed that although the D1R agonist SKF 81297 initially appeared to be the weakest activator of nuclear PKA and ERK1/2, this was driven by a lower percentage of responding cells, not by a lower magnitude of response within individual cells (**Fig. 1F-G**). Importantly, including all cells in the calculation of mean responses produced a different rank-order of drug effects compared to when only the drug-responsive cells were considered (**Fig. 1H-I**). Hence, the proportion of cells that respond to a drug may be an important consideration when comparing the ability of different drugs to activate a given pathway. This granularity would be lost in techniques that rely on the “bulk” analysis of thousands of cells including western blot or plate-reader based assays, both of which are commonly used to assess signaling.

High content imaging approaches can produce rich datasets of secondary measures (e.g. cell morphology) which can complement the analysis of a primary measure (in this case FRET). In **Fig. 2**, we explored how these secondary measures can be used to refine and expand the analysis of the primary measure. We first compared the size and shape (roundness) of nuclei transduced with our FRET reporters to the basal FRET ratio of each cell (calculated as above) and found that basal FRET ratios were evenly distributed across cells with varying size and roundness (**Fig. 2A, D**). We observed that the nuclei of our cultured neurons exhibited a range of distinct nuclear shapes, and given published evidence that MSNs can be distinguished from other striatal cell types based on distinct nuclear morphology ^24^, we hypothesized that cells with 50-125 μm^2^ round nuclei (roundness score >0.9) (**Fig. 2A,D** purple square) would be enriched for MSNs. We assigned cells to four categories based on their nuclear morphology **(Fig. 2A,D** colored squares). Cells in the “medium-round” group were hypothesized to be enriched for MSNs, and these comprised the majority of all cells in culture. We compared the FRET response of each cell plotted against the CFP intensity (a surrogate measure for the expression level of the biosensor) and observed that while FRET responses appeared independent of expression level, cells of this “medium round” morphology tended to exhibit larger FRET responses (data are merged across all treatments) (**Fig. 2B,E**). Within these “medium round” cells we found no association between the expression level of the biosensor and the response magnitude to stimulation (**Fig. 2 C, F**), suggesting that differences in biosensor expression levels do not result in differences in response magnitude.

To further explore our hypothesis that the “medium round” cell group was enriched in striatal MSNs, we again performed high-content imaging, this time on paraformaldehyde-fixed striatal neurons using immunofluorescence to probe for tubulin and the striatal cell marker DARPP-32. Hoechst dye was used to stain nuclei and determine nuclear morphology (**Fig. 3A**). We plotted relative DARPP-32 expression levels on a coordinate plot of nuclear size and roundness, and found that cells with high levels of DARPP-32 expression tended to concentrate in the “medium round” cell grouping (**Fig. 3B**). Re-analysis of the FRET data presented in Figure 1 using these morphological criteria showed that, in terms of both nuclear PKA (AKAR-NLS) and nuclear ERK1/2 (EKAR-NLS) signaling, the response to drug treatments significantly differed between cell-types (2-way ANOVA, Treatment x Cell-Type interaction, p<0.0001, p<0.001 AKAR-NLS and EKAR-NLS respectively) (**Fig. 3C, D**). Specifically, “medium round” cells were more responsive to all drugs in both PKA and ERK1/2 signaling, with several of these comparisons being individually significant. Finally, as an independent method of verifying these differences in drug response, we performed cFos immunofluorescence on striatal cultures stimulated for 3 hours with each drug (**Fig. 3E**). Here we again found that cFos induction significantly differed between cell types (2-way ANOVA, Treatment x Cell-Type interaction p<0.0001) and that relative to all other cell groups, “medium round” cells were significantly more responsive to SKF 81297, NMDA and forskolin stimulation.

**Figure 3:**
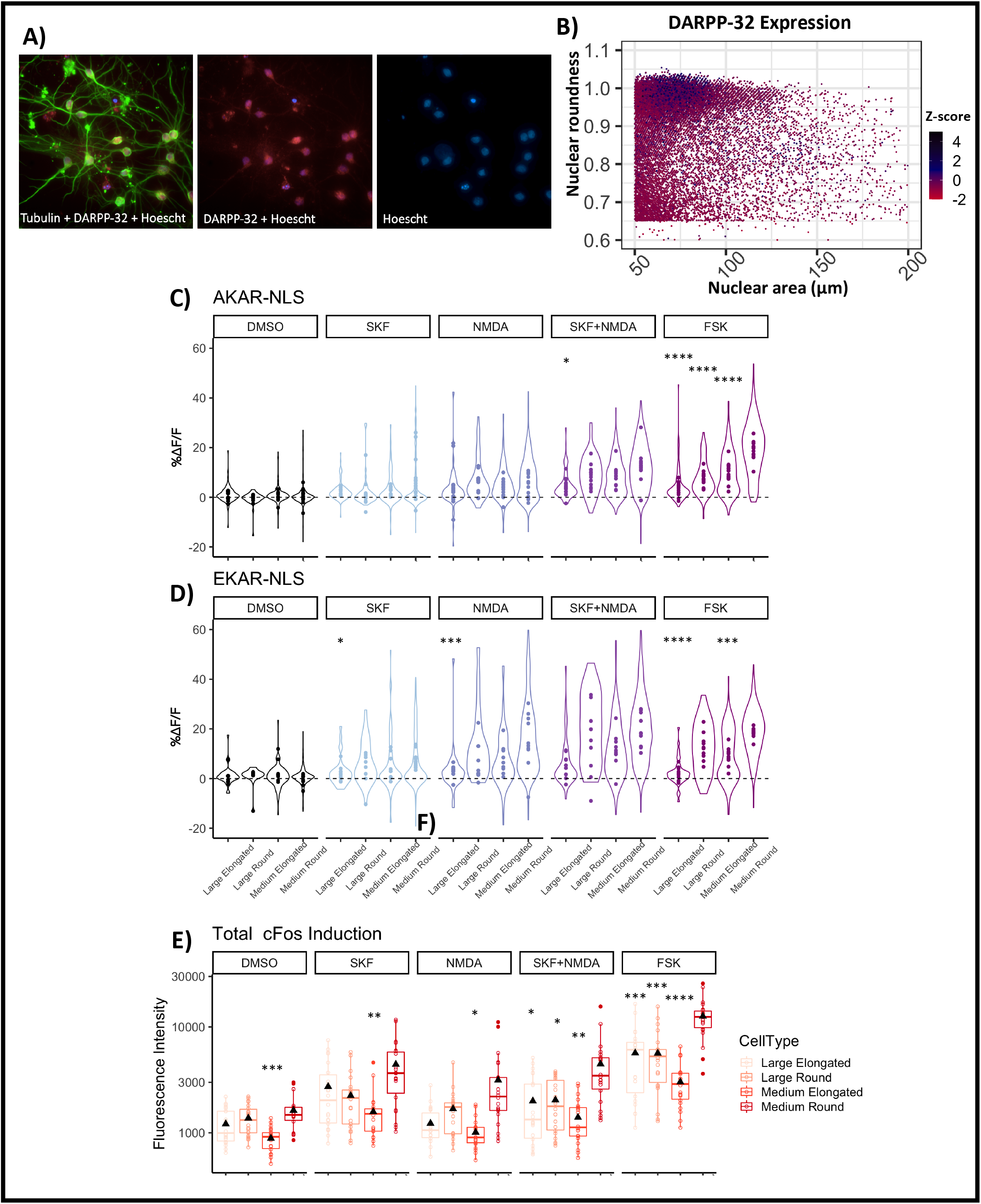
Comparison of cellular and signaling phenotypes of nuclear morphology-defined cell types. A) Immunofluorescent labelling of primary striatal neurons for tubulin and DARPP-32 with Hoechst staining. B) Expression level z-score of DARPP-32 (coded by color) in single cells plotted on a coordinate map depicting measurements of nuclear size and shape. C) Nuclear PKA activation 10 min after indicated drug treatments. D) Nuclear ERK1/2 activation 10 min after indicated drug treatments. Violin plots represent single cell distribution while points represent biological replicates. E) Induction of cFos 3 hours after the start of drug treatments. Statistical comparisons were performed by 2-way ANOVA followed by Bonferroni corrected t-tests comparing “medium round” cells to all other cell types for each treatment. *p<0.05, **p<0.01, ***p<0.001, ****p<0.0001 versus “medium round” cells. Experiments consisted of 3, 10 and 8 biological replicates for DARPP-32, AKAR-NLS and EKAR-NLS, respectively. cFos staining was performed on the cells used in all FRET experiments (n=18). Abbreviations and drug concentrations: DMSO, dimethyl-sulfoxide; SKF, SKF 81297 (1 μM); NMDA, N-methyl-D-aspartic acid (5 μM); and FSK, forskolin (5 μM).

The results presented thus far have focused on *nuclear* signaling responses, using biosensors localized to the nuclear compartment. We also performed a similar analysis of PKA and ERK1/2 signaling in the *cytosol* using biosensors instead tagged with a nuclear export sequence (NES) (**Fig. 4A**). Automated single-cell analysis of images such as that presented in **Fig. 4A** presented an additional challenge due to the presence of long neuronal projections which overlapped with the projections of adjacent neurons. A partial solution to this conundrum was to include only the cell bodies and discard all projections from analysis, as the cell bodies can be easily identified based on their shape. This avoids the potential bias that could be introduced by including multiple “objects” which in fact correspond to different pieces of a single cell. We used this approach to measure cell-body signaling and analyzed the responses as described above (**Fig. 4 B-G**). Compared to nuclear responses, cytosolic signaling responses to the same drugs appeared less pronounced, with fewer high-responding cells. We also observed that whereas nuclear responses occurred almost exclusively in the positive direction, in the cell bodies many cells exhibited pronounced decreases in FRET, particularly for ERK1/2. This may indicate the presence of some basal ERK1/2 activity that is sensitive either to vehicle treatment or to environmental change (i.e. transfer of the plate from incubator to imager).

**Figure 4:**
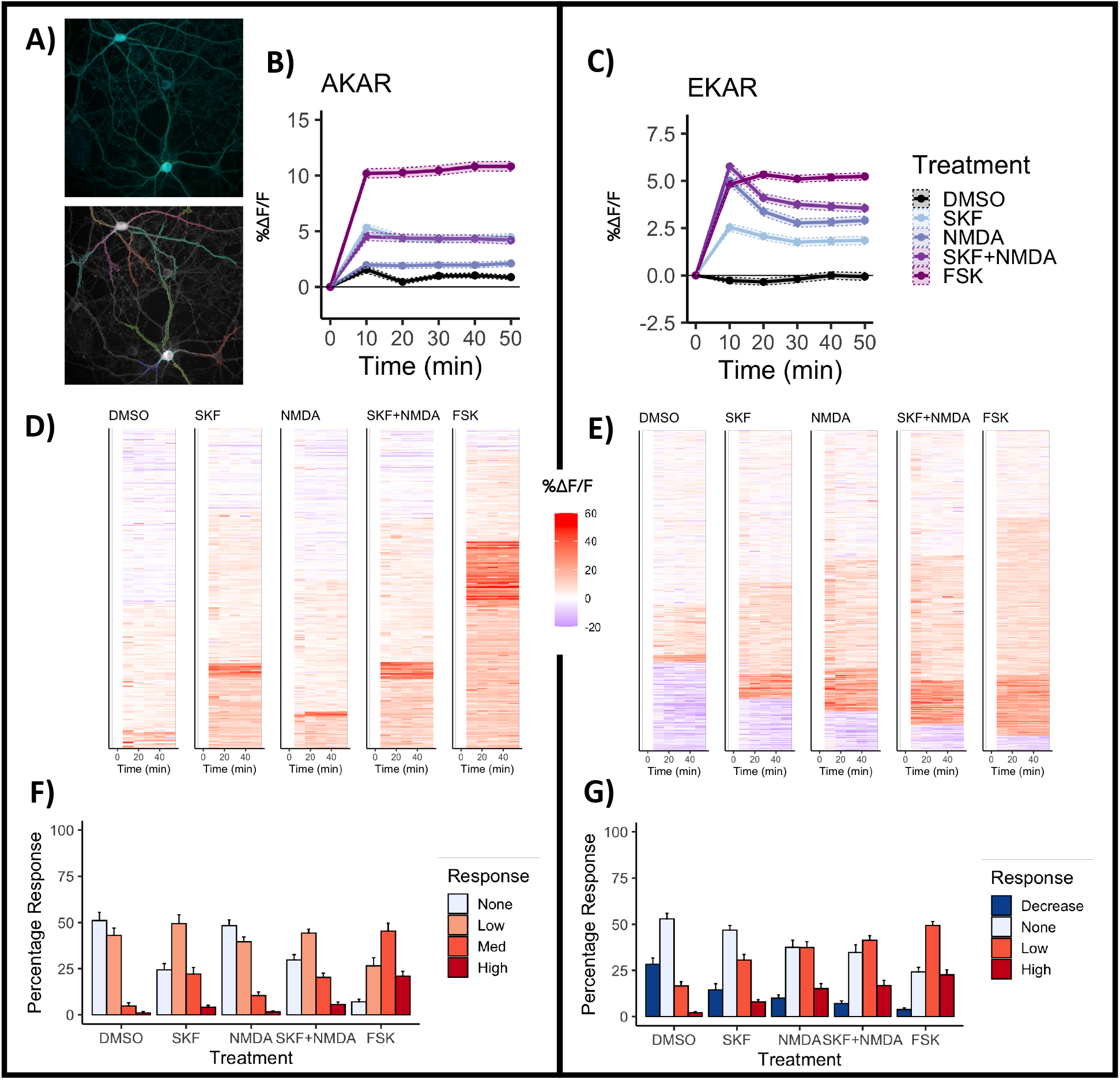
High-content imaging of cytosolic PKA and ERK1/2 activity with FRET biosensors. A) Representative image of striatal neurons expressing a FRET reporter tagged with nuclear export signal (top) and outlines showing object identification B, C) Percent change in FRET ratio (%ΔF/F) averaged across all cells at 10 minute intervals after drug stimulation. Shaded region shows the SEM calculated across cells. D, E) Heatmaps depicting individual cells imaged at 10-min intervals. Cells are organized into four response clusters as described in the following panels. F, G) Percentage of cells falling into each response cluster (mean±SEM across biological replicates). Experiments consisted of 4 biological replicates. Abbreviations and drug concentrations: DMSO, dimethyl-sulfoxide; SKF, SKF 81297 (1 μM); NMDA, N-methyl-D-aspartic acid (5 μM); and FSK, forskolin (5 μM).

In each of our single-cell analyses, we observed intercellular variability not only in the magnitude, but also in the temporal pattern of signaling pathway activation. In **Fig. 5** we explored the analysis of response kinetics by examining the effects of ERK pathway antagonists on nuclear signaling responses. We observed robust activation of ERK1/2 in the nucleus following either D1R or adenylate cyclase stimulation. Activation of ERK1/2 by either D1R or cAMP signaling is thought to occur downstream of PKA and DARPP-32 activation, through several previously described mechanisms ^14–16^. In contrast, the role of ERK1/2 signaling in regulating PKA activity, has not been previously described. To explore this, we measured nuclear PKA and ERK1/2 signaling in the presence of two inhibitors of the ERK pathway: the MEK inhibitor U0126 and the ERK1/2 inhibitor SCH 772984. As expected, both drugs completely abolished the activation of nuclear ERK1/2 (**Fig. 5A**), but we also found that MEK inhibition with U0126 decreases nuclear PKA activation after a 10 minute treatment with D1R agonist or forskolin (**Fig. 5B**). A similar trend was observed when ERK1/2 was inhibited with SCH 772984 (**Fig. 5B**). When we examined the time-resolved single-cell responses, we observed that this inhibitory effect appeared to be transient, with the most pronounced inhibitory effect occurring 10 min after treatment. Single cell responses were visualized by first filtering out non-responding cells (~70% of cells for SKF 81297 and ~10% of cells for forskolin) and then performing kinetic clustering (see Methods), categorizing cells based on the time-course of their response over time (**Fig. 5C-E**). Mean profiles of the four predominant response types identified by clustering are shown in **Fig. 5F**. Thus, when looking at the full time-course it appears that although ERK pathway inhibitors do not diminish PKA activation *per se*, they do delay the nuclear PKA signal, as demonstrated by the reduced number of “fast” responding cells (**Fig. 5G**). In contrast, U0126 had no effect on the activation of PKA in the cell body 10 min after treatment **(Fig. 5H)** suggesting that this effect cannot be explained by non-specific PKA inhibition, and that ERK1/2 may have a specific potentiating effect on PKA activation in the nucleus.

**Figure 5:**
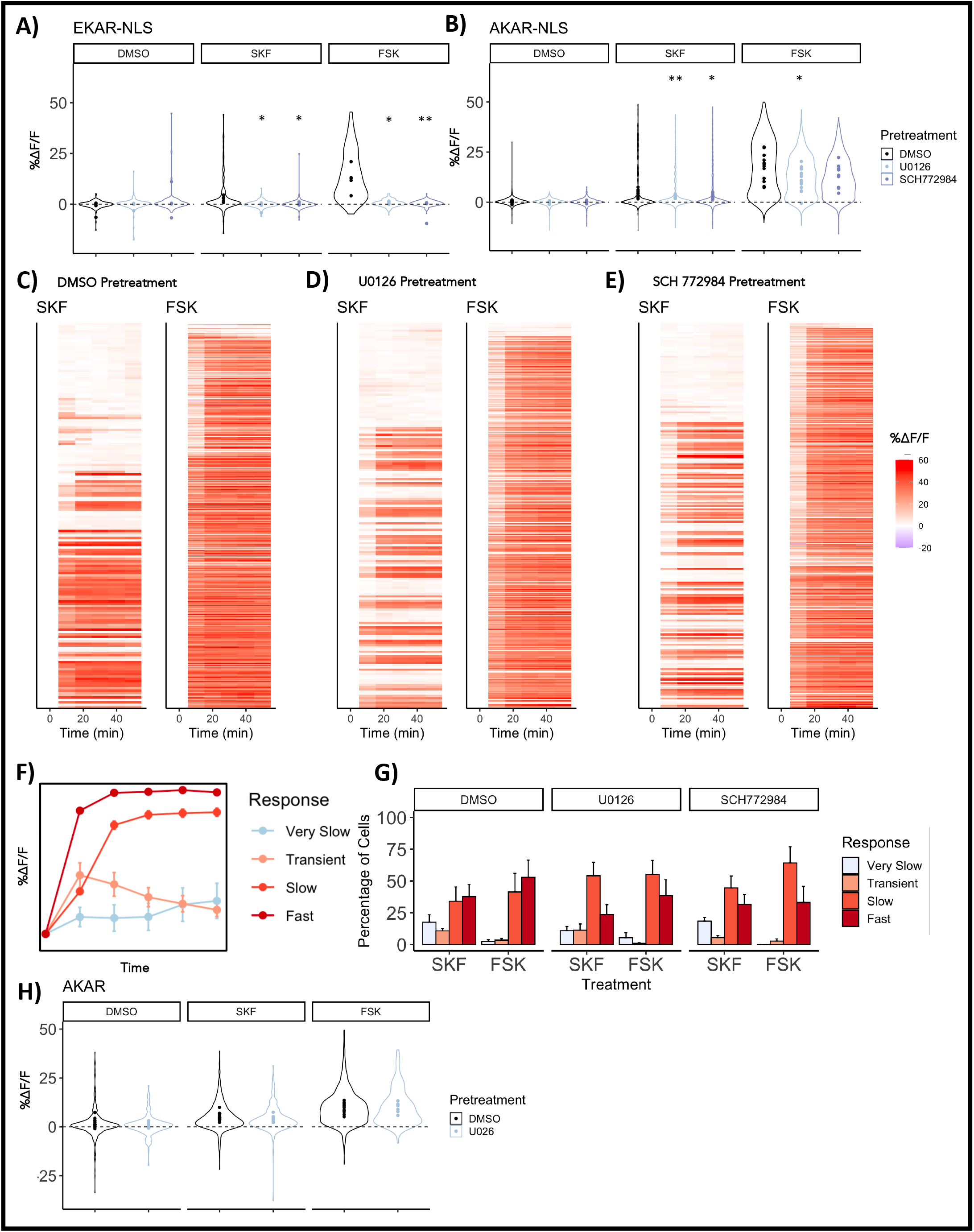
Effect of ERK1/2 inhibition on the kinetics of nuclear PKA activation. A,B) Percent change in FRET ratio (%ΔF/F) for EKAR-NLS and AKAR-NLS, 10 min after stimulation of cells with either SKF 81297 or forskolin, in the presence of the MEK inhibitor U0126 (1 μM), ERK1/2 inhibitor SCH 772984 (10 nM), or vehicle (DMSO). Violin plots show the range of individual cell responses, while points indicate the mean of each biological replicate. C-E) Heatmaps depicting individual cell responses over time under each pretreatment condition. Cells were first filtered to remove non-responding cells, and then clustered based on response kinetics. F) Mean±SEM response of cells in 4 kinetically-defined clusters. G) Percentage of cells falling into each response category across treatment combinations. Experiments consisted of 4 and 8 biological replicates for EKAR-NLS and AKAR-NLS respectively. Statistical comparisons were performed by 2-way ANOVA followed by Bonferroni corrected t-tests comparing each antagonist to DMSO within a given treatment condition. *p<0.05, **p<0.01, relative to DMSO.

## Discussion

Neurons can be categorized into specific types based on many factors, including anatomical location, morphological properties or the expression of specific molecular markers. Classically, cells of a given “type” have often been treated as if they form a homogeneous population. This assumption is typically made out of practical necessity. However, the advent of single cell sequencing and technologies for large scale electrophysiological recording have begun to reveal the extent of intercellular variability within broad cell-type classifications ^25,26^. On the other hand, the most widely used techniques for assaying cellular signaling - notably western blotting or plate reader assays - lack cellular resolution. The same “homogeneity assumption” also predominates in pharmacological studies, where drugs are often described as having a single, quantifiable effect on a cell population of interest. Hence, the possibility of cellular heterogeneity in signaling pathway engagement has traditionally been underrecognized and underappreciated. Standard fluorescent microscopy techniques can resolve individual cells, but lack the throughput for large scale experiments. The methods and results presented here illustrate a simple approach to the application of live-cell biosensors using high-content microscopy that can capture the signaling dynamics of thousands of individual cells grown in multi-well formats.

As the present study shows, individual neurons differ markedly in their response to a given drug. Even forskolin, a drug commonly used as a positive control in studies of GPCR-mediated cAMP signaling, was seen to exert variable effects. In our experiments, the magnitude of PKA activation by forskolin differed by 6-fold between low and high-responding cells, and ~10% of neurons did not respond at all within the timeframe measured. Similar variability was observed with D1R or NMDA receptor activation. The molecular features that give rise to this variability remain to be determined, but could potentially arise from differences in receptor density, or the wiring of intracellular signaling cascades. Analyzing single cell responses also allows for consideration of the *proportion* of responding cells when trying to understand drug effects. This would be impossible using standard approaches that lack cellular resolution, and may obscure important biological differences. For example, seemingly identical results would be produced by a small number of high-responding cells or a large number of weakly responding cells.

Primary neuronal cultures often contain both neurons and glia. In our experiments, we utilized AAVs possessing a neuron-specific promoter to limit the expression of sensors to neurons. This cell-type specific genetic labelling could be further refined using either transgenic animals or viruses to express tools such as Cre and Flp recombinases in cells defined by multiple molecular features ^27,28^. This approach would also allow multiple biosensors to be expressed in distinct neuronal populations, allowing multiple biochemical activities to be tracked simultaneously, in distinct cell types.

A significant limitation of the approach presented here is the limited temporal resolution of high content imaging. The acquisition of many images from multiple wells necessarily takes time, and thus this approach is well suited for detecting signaling processes on a scale of minutes, but not seconds. The desired temporal resolution is an important factor to consider in the design of experiments, as there is a direct tradeoff between the speed of imaging and the number of images acquired, or the number of wells imaged.

In conclusion, high content microscopy, when combined with fluorescent biosensors, can provide a novel vantage point from which to capture the fine-grained nuances associated with cellular signaling dynamics. Although the experiments presented here were conducted on primary rat striatal neurons, we believe the analytic principles can be easily adapted for use on any cell type, with a variety of fluorescent biosensors and imaging hardware.

## Acknowledgements

This work was supported by grants from the Weston Brain Institute and CIHR. J.J.T was supported by doctoral studentships from the CIHR and the McGill Healthy Brains for Healthy Lives initiative. R.M. was supported by studentships from the McGill-CIHR Drug Development Training Program and the McGill Faculty of Medicine. We thank Dr. Michiyuki Matsuda (Kyoto University) for providing us with biosensor constructs and Dr. Hiroyuki Hioki (Kyoto University) for providing AAV plasmids. We thank the McGill Pharmacology and Therapeutics Imaging and Molecular Biology platform, as well as the McGill Advanced Bioimaging facility for training and assistance with microscopy and image analysis. Lastly, we thank members of the Hébert, Clarke and Tanny labs for feedback and guidance throughout the development of the project, and critical reading of the manuscript.

